# Novel loading protocol combines highly efficient encapsulation of exogenous therapeutic toxin with preservation of extracellular vesicles properties, uptake and cargo activity

**DOI:** 10.1101/2023.11.12.566761

**Authors:** Stefania Zuppone, Natasa Zarovni, Kosuke Noguchi, Francesca Loria, Carlo Morasso, Andres Lohmus, Ikuhiko Nakase, Riccardo Vago

## Abstract

Extracellular vesicles (EVs) have mostly been investigated as carriers of biological therapeutics such as proteins and RNA. Nevertheless, small-molecule drugs of natural or synthetic origin have also been loaded into EVs, resulting in an improvement of their therapeutic properties. A few methods have been employed for EV cargo loading, but poor yield and drastic modifications of vesicles remain unsolved challenges. We tested a different strategy based on temporary pH alteration through incubation of EVs with alkaline sodium carbonate, which resulted in conspicuous exogenous molecule incorporation. In-depth characterization showed that vesicle size, morphology, composition, and uptake were not affected. Our method was more efficient than gold-standard electroporation, particularly for a potential therapeutic toxin: the plant Ribosome Inactivating Protein saporin. The encapsulated saporin resulted protected from degradation, and was efficiently conveyed to receiving cancer cells and triggered cell death. EV-delivered saporin was more cytotoxic compared to the free toxin. This approach allows both the structural preservation of vesicle properties and the transfer of protected cargo in the context of drug delivery.

## Introduction

Recently, increasing efforts have been made to exploit nanovectors for drug delivery. They represent a privileged system that offers several advantages over the use of drugs as such. In fact, as the drug is “protected” inside the carrier, it is not subjected to degradation by serum components; therefore, encapsulation often enhances its solubility and enables higher accumulation in target cells. In this context, naturally secreted extracellular vesicles (EVs), which have the intrinsic ability to carry and deliver diverse bioactive cargoes, appear to be a preferable choice compared to synthetic particles, mostly due to the lack of immunogenicity and innate stability in biological fluids. In particular, exosomes represent a class of EVs that have attracted a special attention because of their nanoscale size (30–150 nm), which allows them to easily penetrate almost all body districts and cross biological barriers, as well as their cargo protection and customizable surface, making them ideal vectors for targeted drug delivery in medical applications.^1,2^ They originate from inward budding of the late endosome membrane, which results in the formation of small vesicles inside the lumen. The neo-formed structures, called multivesicular bodies, can either fuse with lysosomes for degradation or with the plasma membrane of the cell, releasing exosomes in the extracellular environment. The molecular composition of thus formed vesicles strongly depends on the cell of origin, containing different biomolecules, such as proteins, carbohydrates, nucleic acids, and lipids, which are finely sorted and transferred to the receiving cells to mediate intercellular communication and regulate many biological processes.^2^ Although we base our attachment to the term “exosome” to more than just a vesicle size (for instance we consider the presence of typical ESCRT proteins from the exosome biogenesis pathway), in congruence with the latest suggestions on EV nomenclature^3^ we refer to exosomes as nano-sized EVs (nsEVs) throughout this article.

To date, multiple research groups have been developing an efficient approach to incorporate bioactive molecules into nsEVs. The use of active loading methods that exploit mechanical (extrusion), chemical (detergent permeabilization), or physical stimuli (sonication, electroporation, freeze and thaw cycles), for instance, has yielded positive results in terms of increased encapsulation efficiency. However, the major drawback of these techniques is that the structural architecture of vesicles is critically affected, often triggering vesicle aggregation and alteration in size.^4–11^ Electroporation, in particular, has been widely used to incorporate a broad spectrum of druggable molecules into EVs, including nucleic acids such as DNA,^12^ siRNA,^13^ and miRNA,^14^ proteins,^6,8,15^ and small molecules,^16,17^, claiming to have achieved increased loading efficiency compared to other methods. Noteworthy, its loading efficiency varied consistently in different studies,^18^, fundamentally depending on type of exogenous cargo; it was typically reported high for small RNAs and drugs (0.5–60%) and very low for proteins (<0.6%). Consequently, fine optimization of the electroporation protocol is required taking into consideration the properties of the drug used, in order to allow efficient particle engineering, whereas the efficacy of the resulting complexes is significantly hampered by molecular aggregation and structural damage of the vesicles.^19^ The passive loading strategies, modeled on the incubation of EVs/cells with free drugs, are simple and do not compromise the vesicles membrane integrity but, on the other hand, often lead to a very low cargo-incorporation,^9,11,18^ with active loading methods being generally considered superior.

To overcome these challenges, in the present study, we tested the efficiency of a new strategy based on the use of a highly concentrated alkaline buffer to induce conspicuous incorporation of exogenous molecules into nsEVs with a higher loading efficiency compared to electroporation, while preserving the structural and functional features of the vesicles. In this study, we applied our protocol to hydrophilic molecules, with the ultimate goal of loading into EVs a small protein proposed as an anticancer agent. Encapsulation was shown to protect the cargo from enzymatic degradation and to favor the efficient delivery of a toxic payload, which promotes the death of recipient cancer cells. Altogether, we designed a simple protocol to efficiently load nsEVs with functional exogenous therapeutic molecules and preserve their integrity and uptake.

## Material and methods

### Cell culture

HEK293 (human embryonic kidney-derived) cells were purchased from ATCC. HeLa (human cervical cancer-derived) cells were purchased from the Riken BRC Cell Bank (Ibaraki, Japan). 5637 human bladder cancer cell line was purchased from Cell Lines Service. Cells were cultured in DMEM (Gibco, Life Technologies Corporation, Grand Island, NY, USA) (HeLa and HEK) of RPMI (5637) + Glutamax supplemented with 10% FBS and antibiotics (100 U/mL penicillin and 100 μg/mL streptomycine-sulphate). Cells were grown in incubation at 37 °C in 5% CO_2_.

### NsEV isolation

Human embryonic kidney cell line HEK293T (5 × 10^6^) was seeded in 150 mm dishes in DMEM medium supplemented with EV-depleted 10% FCS, 2 mM L-Glutamine, and antibiotics. After 72 h of incubation, the cell culture medium was collected and centrifuged at 300 g for 25 min to remove cell debris. For nsEV isolation, the obtained supernatant was filtered through a 0.22 μm filter before ultracentrifugation at 150,000 g for 2 h at 4 °C (Beckman Coulter). The nsEV-containing pellet was resuspended in phosphate buffer saline (PBS).

### Nanoparticle tracking analysis (NTA)

Nanoparticle tracking analysis (NTA) was performed on the isolated vesicles using a NanoSight LM10-HS microscope (NanoSight Ltd., Amesbury, UK) as previously described.^34^ The NanoSight system was calibrated by using polystyrene latex microbeads. The videos were analyzed using NTA software version 2.3 to determine the concentration and size of the measured particles with the corresponding standard error (SEM).

### Transmission electron microscopy (TEM) analysis

Freshly purified nsEVs from culture media of CD9/CD63-RFP stably expressing HEK293 were absorbed on glow discharged carbon coated formvar copper grids, washed with water, contrasted with 2% uranyl acetate, and air-dried. The grids were observed under a Zeiss LEO 512 transmission electron microscope (Zeiss). Images were acquired using a 2k × 2k bottom-mounted slow-scan Proscan camera, controlled by EsivisionPro 3.2 software.

### Total EV protein quantification

The amount of proteins was quantified for each sample and time point using Pierce™ Bicinchoninic Acid (BCA) Protein Assay (Thermo Fisher Scientific). Samples were loaded at 10 µL per well in duplicate, and the protein content was determined by alignment with the standard calibration curve provided by the kit.

### Lectin Binding Assay

Ten microliters of HEK nsEVs from PBS and carbonate buffer treatments, corresponding to 10^9^ particles, were loaded into flat-bottom black plates with a high binding capacity (Biomat, Cat. No.: MGB01F2-HB8) and diluted with PBS to a final volume of 50 µL. After overnight incubation at room temperature, the plates were washed three times with 300 µL of PBS + 0.05% Tween20 washing buffer, and 50 µL (20 µg/mL) of ConA-FITC and WGA-FITC (Vector Laboratories) were loaded into each well. The 30 min incubation with fluorescently-labelled lectins was carried out at room temperature, after which the washing step with PBS + 0.05% Tween20 was repeated, and 200 µL of PBS was added to each well for measurement of the fluorescence intensity (excitation 485 nm / emission 535 nm) on microplate reader GENios Pro (Tecan).

### Enzyme Linked ImmunoAssay (ELISA) for EV tetraspanins quantification

The surface marker expression of common EV tetraspanins (CD63 and CD9) was assessed using Exo-TEST, an in-house sandwich enzyme linked immunoassay (ELISA) kit (HansaBioMed Life Sciences), following the manufacturer’s protocol provided with the kit. After incubation with PBS or carbonate buffer, 10^9^ HEK nsEVs were loaded onto anti-CD63 coated plate and incubated overnight at 37 °C. Subsequently, biotinylated anti-CD9 antibody (diluted 1/500) included in the kit was used for the detection of bound EVs in combination with streptavidin-HRP (diluted 1/5000). The absorbance at 450 nm was measured on microplate by a GENios Pro microplate reader (Tecan).

### Cholesterol Assay

The lipid content of HEK nsEVs after incubation with PBS or carbonate buffer was quantified using a Cholesterol assay (Hansabiomed Life Sciences, Estonia). For this purpose, 10^9^ particles were loaded onto black flat-bottom plates with no binding capacity (Biomat,Cat. No.: MGB03F2-NB) and diluted with PBS to a final volume of 20 µL. The reaction mix preparation, incubation, and fluorescence intensity measurements (excitation 540 nm/emission 590 nm) were performed according to the manufacturer’s guidelines.

### Nucleic acid qualitative and semi-quantitative assessment

Qualitative and semi-quantitative assessment of nucleic acid content was performed using a fluorescence microplate assay, a black 96-well microplate with no binding capacity[(HansaBioMed Life Sciences) and Quant-iT™ RiboGreen™ RNA reagent (Invitrogen; Cat. R11491; 1/1000 working dilution). Briefly, samples, 10X diluted in DPBS (Sigma-Aldrich; Cat.: D8537-500ML) to a final volume of 50 µL per well, were loaded onto microplate wells, followed by incubation with 50 µL per well of dye stock solution (1/500 dilution in DPBS) for 30 min at RT. Fluorescence intensity (λex = 485 nm, λem = 535 nm, gain:50) was measured using a Tecan GENios Pro microplate reader. The results were reported as signal-to-background ratios.

### Fluorescent labelling of nsEVs and *in vitro* uptake tracing

NsEVs isolated from the cell culture media were resuspended in PBS and combined with a PBS solution containing Vybrant^TM^ DiO (1 μl for 100 μg nsEVs, Molecular Probes, Eugene, OR, USA). The samples were incubated at 37 °C for 30 min (protected from light) under stirring and then passed through Exosome spin columns (MW 3000, Invitrogen) to remove any incorporated dye. PBS containing the dye alone was used as a negative control. The samples were then ultracentrifuged (150,000 × g, 2 h at 4 °C) and the pellet was resuspended in PBS. For the *in vitro* tracing assay, 5637 were seeded (up to 50–60% confluence) in a 24 multiwell plate and incubated at 37 °C with 5% CO_2_. 24 hours later, labelled nsEVs (5 μg or 10 μg/well) were added to recipient cells. The uptake of the NsEVs was evaluated after 24 hours. Fluorescent images were captured using Axio Vision Imaging Software (Axiovision Rel 4.8®) on an Axio Imager M2 microscope (Carl Zeiss, Oberkochen, Germany).

### Sodium carbonate-based molecule encapsulation into isolated nsEVs

HEK293 derived-nsEVs were resuspended with dextran-FITC (Sigma-Aldrich) or seed saporin (SAP) (Sigma-Aldrich) in 1 M Na_2_CO_3_ (pH 11.5) or PBS and incubated for 30 min at 37 °C. The amount of nsEVs used was calculated as protein concentration. A ratio of 1 μg nsEVs to 1/1.5/2 μg dextran/SAP was used to define the optimal yield of exogenous protein encapsulation. The pH was neutralized by dropwise addition of HCl, and the sample was filtered by centrifugation through Exosome spin columns 100 kDa MWKO (Merck Millipore) to remove any incorporated proteins and washed with PBS or NaCl 150 mM to reduce protein aggregation. Dextran, SAP or nsEVs alone were used as negative controls.

### Encapsulation of fluorescently-labelled dextran and SAP into nsEVs by electroporation

To load fluorescently-labelled dextran or SAP into nsEVs, nsEVs (25 μg) were mixed with FITC-labelled dextran (70 kDa) or SAP (25 kDa) in PBS (100 μl) at a total protein ratio (w/w) of 1:1, 1:1.5, or 1:2. After electroporation (poring pulse: two pulses (5 ms), transfer pulse: five pulses (50 ms)) in a 1 cm electroporation cuvette at room temperature using a super-electroporator NEPA21 Type II (NEPA genes, Tokyo, Japan), the unencapsulated FITC-dextran or FITC-SAP was removed by washing and filtration using Amicon Ultra centrifugal filters (100 K device, Merck Millipore). The incorporation of FITC-dextran and FITC-SAP into the nsEVs was confirmed by spectrofluometer analysis (FP-6200, JASCO, Tokyo, Japan). The incorporation of FITC-dextran and FITC-SAP into vesicles was quantified by interpolation based on a calibration curve generated using scalar dilutions of FITC dextran/SAP. The electroporation method was optimized for poring pulses (50, 100, and 150 V), resulting in the encapsulation of FITC-dextran (15–20 ng/ml in 50 μg/ml nsEVs), respectively. Therefore, we used an experimental poring pulse of 50 V for electroporation.

### Confocal microscopy

HeLa cells (2 × 10^5^ cells, 2 ml) were plated on a 35 mm glass dish (Iwaki, Tokyo, Japan) and incubated in the cell culture medium. After complete adhesion, cells were washed with culture medium and treated for 24 hours with nsEV samples (25 µg/ml), either treated with electroporation or sodium carbonate buffer protocol to load dextran-FITC. Cells were then washed with fresh cell culture medium and analyzed using an FV1200 confocal laser scanning microscope (Olympus, Tokyo, Japan) equipped with a 40x magnification objective.

### Flow cytometry analysis

5637 cells were cultured in 24-well microplates until 60% confluence. 24 hours later, the cells were treated with labelled nsEVs (15 μg/well) and incubated at 37 °C, 5% CO_2_ for 48 hours. The cells were detached by incubation with 0.01% trypsin for 5 min at 37 °C, collected, washed with 1% FBS (in PBS), and subjected to flow cytometric analysis using Accuri^TM^ flow cytometer. Analysis was performed for 20,000 gated events per sample.

### Protein extraction, western blot and silver staining analysis

Cells were washed twice with cold PBS, collected by scraping, centrifuged 5 min at 1200 rpm, and lysed for 30 min in ice-cold buffer (150 mM NaCl, 2 mM NaF, 1 mM EDTA, 1 mM EGTA, 1 mM Na_3_VO_4_, 1 mM PMSF, 75 mU/ml aprotinin-Sigma-, 50 mM Tris-HCl, pH 7.5) containing 1% TX-100, and a cocktail of protease inhibitors (Sigma-Aldrich). Cell lysates were centrifuged at 10,000 × g at 4 °C for 10 min. Proteins contained in the supernatant were quantified by BCA assay, resuspended in sample buffer (62.5 mM Tris-HCl (pH = 6.8), 2% SDS, 10% glycerol, 0.002% bromophenol blue), and boiled at 95 °C for 5 min. The same amount of total protein was loaded unless otherwise indicated. Proteins were then separated *via* SDS-PAGE under non-reducing conditions for the detection of the CD9, CD63, and CD81 or under reducing conditions (with 5% 2-mercaptoethanol) for the detection of the other proteins. Proteins were silver stained or transferred onto a nitrocellulose membrane for western blot analysis incubated with 5% non-fat powdered milk in TBS-T (0.5% Tween-20) for 1 hour and then with the following antibodies: saporin (1:5000), flotillin (1:5000, Sigma), Alix (1:800, Santa Cruz Biotechnology), anti-CD9 (1:10,000; BD Pharmingen), anti-CD63 (1:20,000, BD Pharmingen), anti-CD81 (1:5000, BD Pharmingen), anti-TSG101 (1:1000; Novus Bio). Secondary horseradish peroxidase-conjugated antibodies (anti-mouse/rabbit IgG HRP-linked whole antibody donkey; GE Healthcare) were used, and the immunoreactive bands were visualized using Enhanced Chemiluminescence (Merck Millipore).

### Acquisition of Raman spectra, data processing and analysis

Raman spectra were acquired using an InVia Reflex confocal Raman microscope (Renishaw plc, Wotton-under-Edge, UK) equipped with a solid-state laser light source, operating at 785 nm. We visualized lipids and proteins in the sample by imaging the Raman peaks at 2935 and 2853 cm^−1^ originating from methyl groups in proteins and methylene groups in fatty acids. The intensity of the peak indicates the overall protein and lipid content, whereas a shift in the peak would indicate conformational alterations. The Raman spectrometer was calibrated daily using 520.7 cm^−1^ of a silicon wafer. Typically, a 3 µL drop of sample was dropped onto the surface of Raman-compatible CaF_2_ discs (Crystran, UK) and dried for 20 min at room temperature. The Raman study was performed using a 785 nm excitation laser with 100% power (approximately 42 mW at source), a 1200 l/mm grating and a 100x objective. Spectra were acquired as the sum of nine 10 s acquisitions. The spectral resolution was ~about1 cm^−1^. For each sample, four different spectra were collected at different positions of the drop. The software package WIRE 5 (Renishaw, UK) was used for spectral acquisition and to remove the cosmic rays. Background fluorescence was removed using an asymmetric least-squares smoothing method. ^35^ The spectra acquired for each sample were vector-normalized and averaged. Spectrum normalization and data analysis were performed using OriginPro, Version 2019 (OriginLab Corporation, Northampton, MA, USA).

### Cell viability assay

5637 cells (5×10^3^ cells/well) were seeded in 96 wells plates and incubated for 72 hours with 2, 10, or 20 µg SAP-bearing nsEVs or 0.1, 1, or 10 µg SAP alone (seed SAP). After 72 hours of incubation at 37 °C, 3-(4,5-dimethylthiazol-2-yl)-2,5-diphenyltetrazolium bromide (MTT) (5 mg/ml in PBS) was added (0.5 mg/ml working concentration). After 1 hour incubation at 37 °C, the supernatants were removed, and 100 μl/well of dimethyl sulfoxide was added to dissolve the formazan crystals. Cell viability was assessed by measuring the absorbance at 570 nm and expressed as a percentage of the untreated control. SAP toxicity was evaluated as IC_50_, shown as the mean ±SEM from three independent experiments.

### Statistical analysis

Three biological replicates were performed for all *in vitro* experiments. When appropriate, statistical significance was determined using 2-tailed Student’s *t* tests. Test symbols mean: \*\**p* < 0.01; \*\*\**p* < 0.001.

## Results

### A pH alteration *via* sodium carbonate-based exposition enhances exogenous molecules incorporation and preserves nano-sized EVs structure and uptake properties

Since extracellular vesicles reflect the cell-of-origin molecular composition, it is very important to select the proper cell line to produce safe carriers for therapeutic applications. In this study, we opted for commercially available HEK293 cells as vesicle donors because they have been reported to be one of the most frequently used systems for this purpose, including EV-based therapeutics that are currently in clinical trials.^20^ Besides, HEK293 cells and their subtypes are widely used in the production of biotherapeutics such as recombinant proteins and viruses.^21^

We set up and optimized the protocol for the isolation and purification of EVs from HEK293 culture medium, consisting of sequential centrifugation and ultracentrifugation steps, and a 0.22 μm filtration to enrich for nsEVs.^25^ We then evaluated the impact of incubation in a carbonate buffer on the physiological properties of the collected vesicles. We performed accurate molecular and structural characterization of EVs, as much as possible in accordance to the updated guidelines of the International Society for Extracellular Vesicles (MISEV2018).^22^ After recovery, nsEVs appeared to be perfectly preserved in their morphological and biophysical properties, as well as in their overall number, as shown by TEM and nanoparticle tracking analysis (Figure 1A and B). Independent of the incubation of isolated EVs in PBS or carbonate buffer, TEM analysis revealed vesicles to be homogeneous in shape, with a size distribution picking at 150 nm in diameter, as determined also by NTA. Typical EV markers (CD63 and CD9) and exosome markers (Alix and TSG101) were still present in nsEVs after treatment (Figure 1C). The slight reduction in these proteins observed in WB was in line with silver staining, which showed a general but modest decrease in total proteins (Figure 1D). Accordingly, in the Raman spectra, the peak at 2930 cm^−1^, relative to CH_2_ in proteins, appeared to be slightly reduced,^23^ confirming that treatment with sodium carbonate affects the protein content of recovered nsEVs, even if not significantly (Figure 1E). At the same time, no shift in peak position was observed that would indicate a change in the protein structure. Notably, we did not observe any variation in the intensity of the peak at 2851 cm^−1^ which corresponds to methyl groups in lipids (Figure 1E).

**Figure 1.**
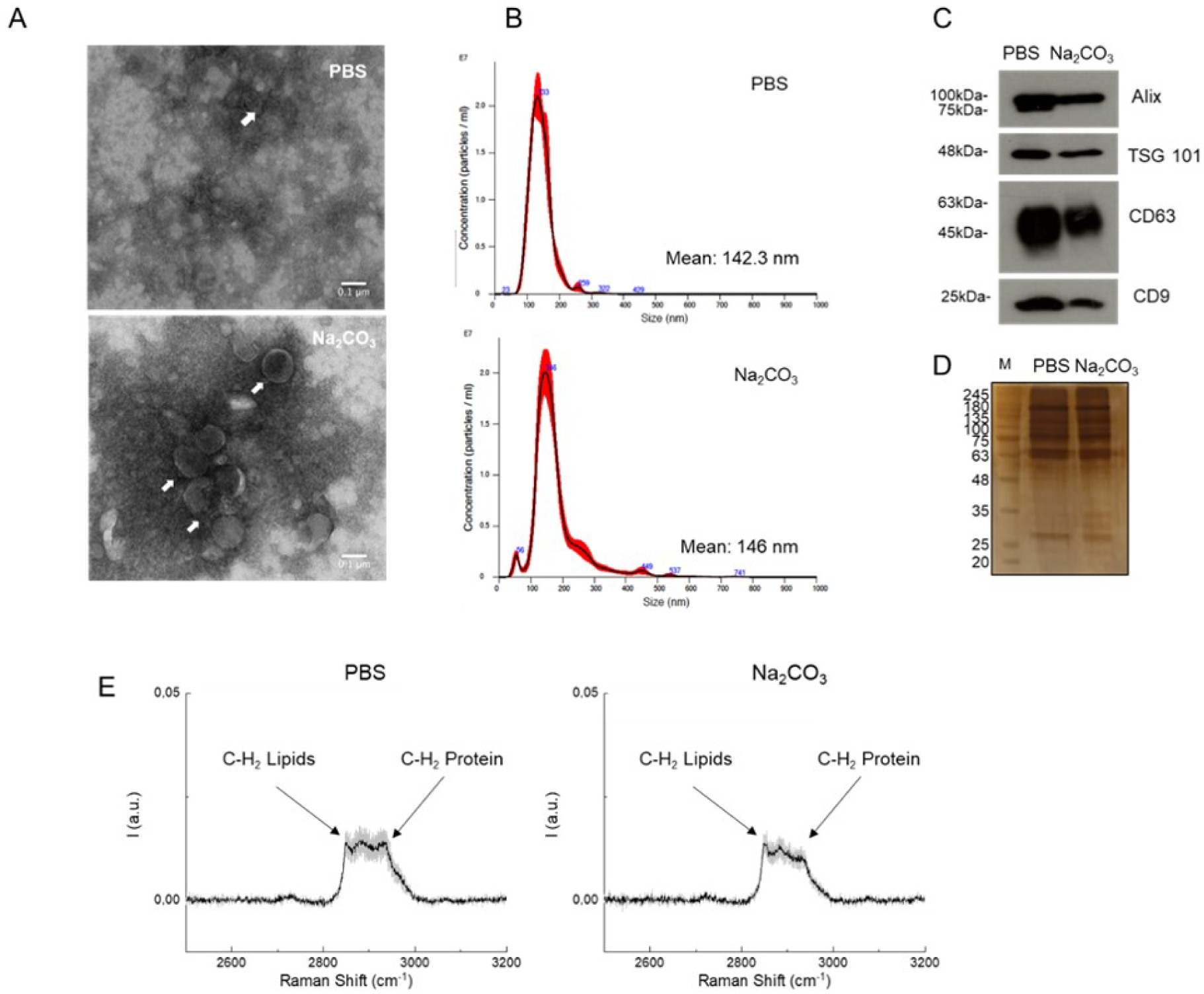
Evaluation of structural modifications of nsEVs after sodium carbonate treatment. Transmission electron microscopy (TEM) image (A) and nanoparticle tracking analysis (B) of HEK293-derived nsEVs after sodium carbonate treatment. Western blot analysis (C) of Alix, TSG101, CD63 and CD9 before (pre) and after (post) sodium carbonate treatment. Comparison of nsEVs treated with phosphate buffer saline (PBS) or sodium carbonate (Na_2_CO_3_) through silver staining (D) and Raman spectroscopy (E). M: molecular weight marker.

These initial results were double-checked and confirmed by additional analytical package assessing, in addition to particle numbers and overall protein content (Figure 2A and B), also the content of cholesterol content, which is known to be enriched in EV membranes from mammalian cells (Figure 2E). We also assessed the preserved immunoreactivity of surface-displayed proteins (CD63 and CD9 in ELISA) (Figure 2C) and surface glycans (by assessing lectin binding) (Figure 2D).

**Figure 2.**
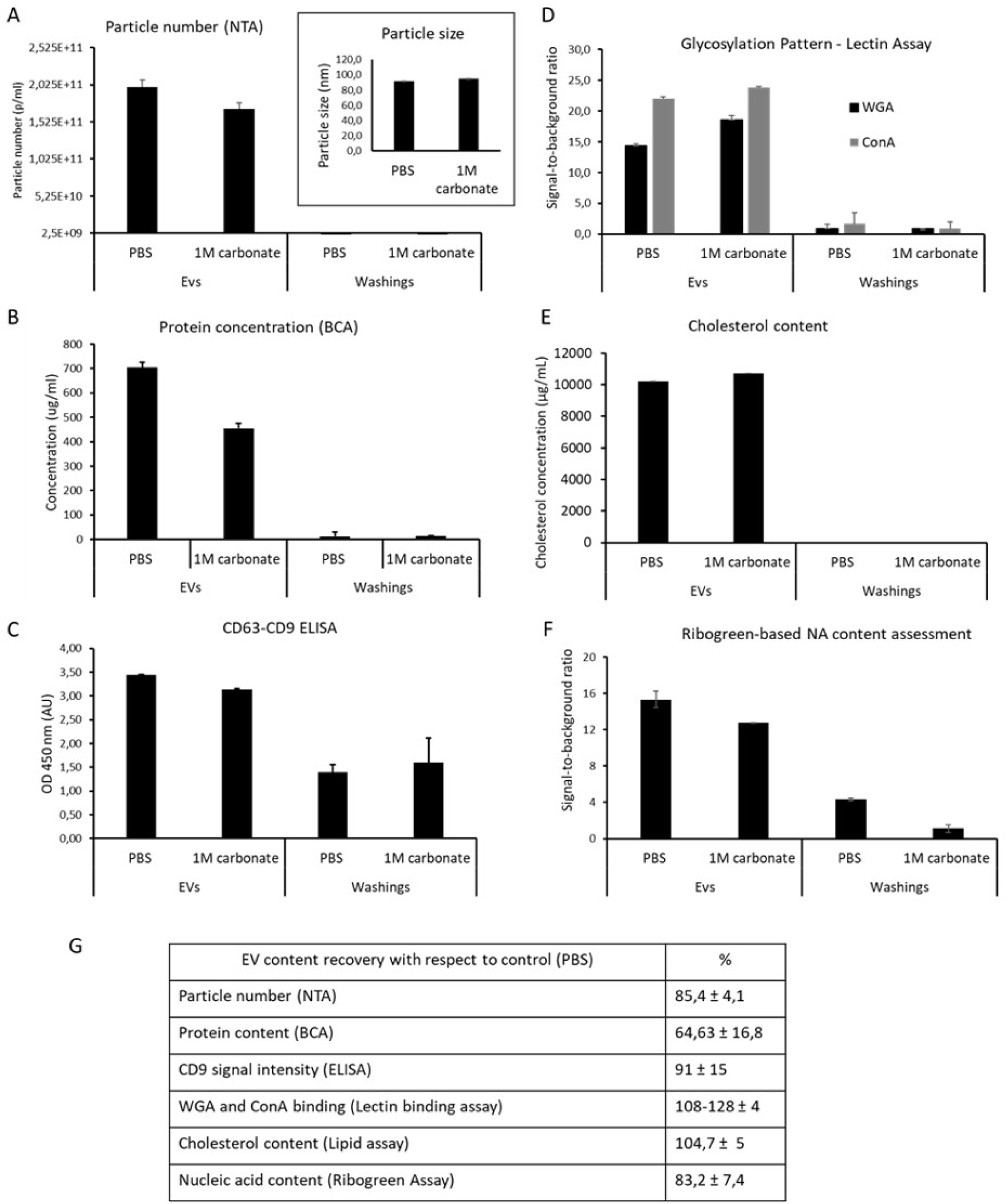
Evaluation of nsEVs content and integrity after sodium carbonate treatment. The number and size of particles recovered upon nsEV incubation in either PBS or in carbonate buffer was assessed by NTA (A). The overall protein concentration was measured by Bicinchoninic Acid Assay (BCA) (B). 10^9^ of particles were loaded per well of 96-well microplate and following assays are performed for the measurement of cholesterol content (E), assessment of surface glycans by lectin (ConA and WGA) binding (D), and quantification of total nucleic acids content via Ribogreen Assay (F). In table G the EV content recovery with respect to PBS is expressed as percentage. All assays are performed according to manufacturer’s indications for commercial kits or otherwise detailed in Material and Methods. The fluorescent signal measured at the plate reader fluorimeter and signal expressed either as a ratio over assay background control (PBS) or as analyte concentration using kit provided standard calibration curve (for cholesterol). Surface tetraspanin expression was assessed by sandwich immunoassay (ELISA) featuring precoated aCD63 microplate and aCD9 detection antibody according to a protocol provided within the kit (see Material and Methods). The absorbance (OD 4500nm) is measured on a bench top microplate reader.

Finally, the overall nucleic acid content of the recovered EVs was assessed using the RiboGreen assay (Figure 2F). Overall results showed a high integrity of vesicular membranes and content, with a decrease in particle number being less pronounced than that of overall proteins (recovery of 85% and 65%, respectively), and in line with maintained nucleic acid content (83%) (Figure 2G).

The drop in particle number was not associated with a change in particle size; therefore, aggregation was unlikely to occur. Moreover, alkaline buffers are known to decrease the rate of aggregation by removing impurities and are therefore used for this purpose in various purification methods. Given the highly preserved cholesterol content, it is possible that the measurement of particle number and protein content overestimates the loss of nsEVs and instead indicates the loss of non-EV contaminants, possibly protein aggregates that can give a signal in NTA, and/or material loosely associated with the EV membrane. Indeed, it is known that the use of a carbonate buffer disrupts the association of peripherally associated proteins or soluble proteins without disrupting the membrane.^24^ It is therefore possible that higher lectin binding (>100% with respect to PBS treated EVs) is due to a cleaner preparation and better exposure of surface glycans.

Finally, to illustrate the preservation of both intrinsic nsEV structure and, possibly, the integrity of engineered membrane proteins upon carbonate buffer exposure, we took advantage of fluorescent vesicles produced in-house from a recombinant HEK293 model expressing the CD9-RFP fusion protein.^25^ As shown in Supplemental Figure 1, western blot analysis demonstrated that incubation in neither PBS nor sodium carbonate altered the CD9-RFP protein structure, as only CD9 associated RFP and no free fluorescent proteins was detectable. Concordantly, recombinant vesicle fluorescence was well appreciated in a fluorimeter (not shown). Therefore, both intrinsic and engineered EV membrane molecules are well-preserved and functional.

### Comparative assessment of nsEV loading efficiency upon alkaline buffer treatment or electroporation

The effect of sodium carbonate treatment on vesicle loading efficiency was compared to that of electroporation as a benchmark. To this end, we initially used FITC-labelled dextran with a molecular size of 70 kDa as a trackable fluorescent payload. This small sugar molecule is commonly used to study the in-cell transport, cell permeation, and encapsulation efficiency. No significant differences in loading capacity in terms of total cargo encapsulation could be observed between the two treatments; notably, the cargo loading efficiency remained low, around 0.2–0.3%. In both cases, the dextran-FITC incorporation capacity was shown to be directly dependent on the amount of payload used when different nsEVs:dextran ratios were used (Figure 3A). The similar performance of the two treatments was further confirmed by the uptake assay performed on HeLa cells, in which a comparable green punctuation was detectable after 24 hours on confocal microscopy (Figure 3B).

**Figure 3.**
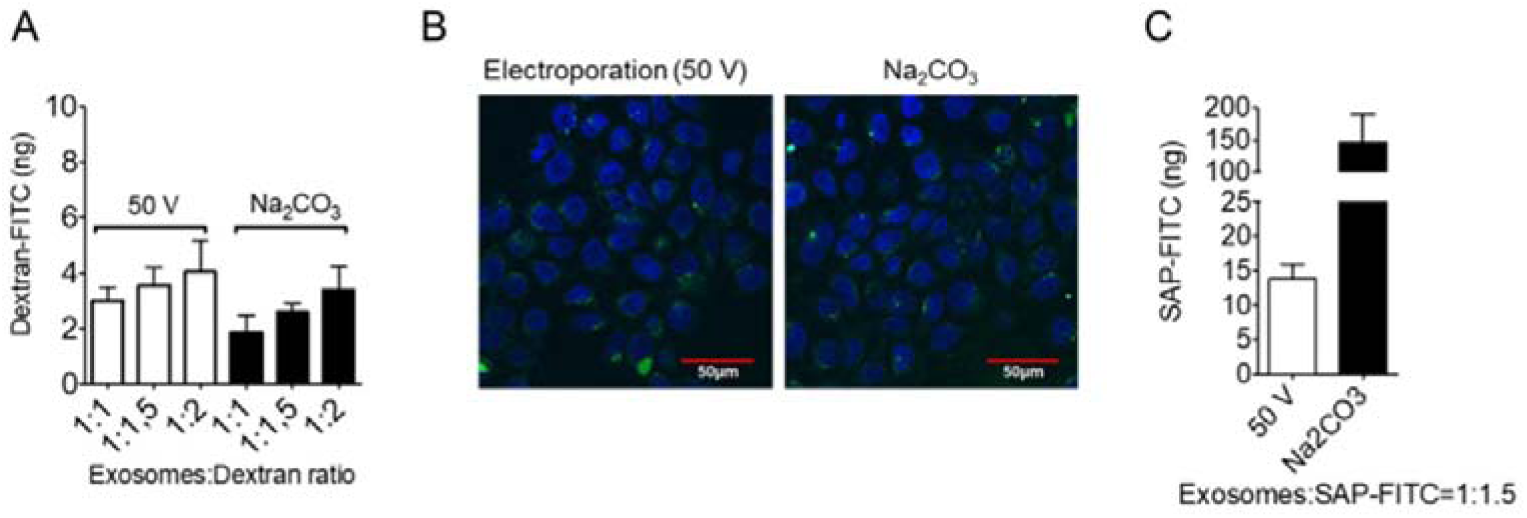
Encapsulation efficiency evaluation and comparison between electroporation and sodium carbonate treatment. A. Spectrofluorometer analysis for quantification of Dextran-FITC incorporation after 50 V electroporation (see material and methods) or sodium carbonate treatment of HEK293-derived nsEVs (25 µg/ml) in various nsEVs:Dextran-FITC ratios (1:1, 1:1.5 and 1:2). B. Confocal microscopy images of Hela cells treated for 24 hours with Dextran-FITC bearing nsEVs (25 µg/ml) after electroporation (50 V) or sodium carbonate incubation in NsEVs:Dextran-FITC=1:1.5 molecular ratio (blue = DAPI; green = Dextran-FITC bearing nsEVs). C. Spectrofluorometer analysis for quantification of SAP-FITC incorporation after 50 V electroporation (see material and methods) or sodium carbonate treatment of HEK293-derived nsEVs (25 µg/ml) in nsEVs:SAP-FITC = 1:1.5 ratio.

Next, we investigated whether sodium carbonate encapsulation efficiency was dependent on the structural properties of the payload. To achieve this goal, we used a 25 kDa plant derived globular protein, saporin (SAP), whose enzymatic activity is directed at inhibiting protein synthesis.^26–29^ First, we used a fluorescently labelled protein, SAP-FITC, and an nsEV-to-cargo ratio of 1:1.5 (w:w). Strikingly, while in the case of electroporation, SAP-FITC incorporation was 4–5 times improved compared to that of dextran, reaching a loading efficiency of 1.5%, sodium carbonate treatment reached a 50 folds higher loading capacity of SAP-FITC with respect to dextran, significantly overperforming the electroporation (Figure 3C) and reaching a loading efficiency of 15%. This demonstrates the relationship between encapsulation efficiency and steric hindrance of the molecule used. The use of higher electroporation voltages (100 V and 150 V) together with higher nsEV:SAP ratios (1:2) seemed to poorly improve SAP-FITC encapsulation, but instead caused vesicle damage, as shown by reduced total protein recovery after treatment (Supplementary Figure 2). Overall, in our settings, the carbonate buffer-based strategy resulted in a 10-fold more efficient loading of a 25 kDa protein into nsEVs with respect to electroporation.

### Sodium carbonate treated-SAP-encapsulating nsEVs exert a high cytotoxic activity on cancer cells

The ultimate way for assessing the success of a loading strategy involves a specific *in vitro* or *in vivo* biological outcome. Hence, we investigated whether SAP loaded nsEVs could deliver and efficiently release their cargo *in vitro* to recipient cells. To answer this question, we first assessed whether sodium carbonate treatment affects cellular uptake. Upon exposure to PBS or carbonate buffer, vesicles were labelled with a lipophilic fluorescent dye and co-incubated with 5637 bladder cancer cells for 24 hours. As shown by microscopy and flow cytometry analysis, no significant differences were observed between the fluorescence signals observed in cells receiving PBS-treated or sodium carbonate-treated nsEVs, further indicating that short-term pH modification did not alter the capacity of the nanovesicles to be captured by cells (Figure 4). As expected, the fluorescence signal depended on the dose of nsEVs used.

**Figure 4.**
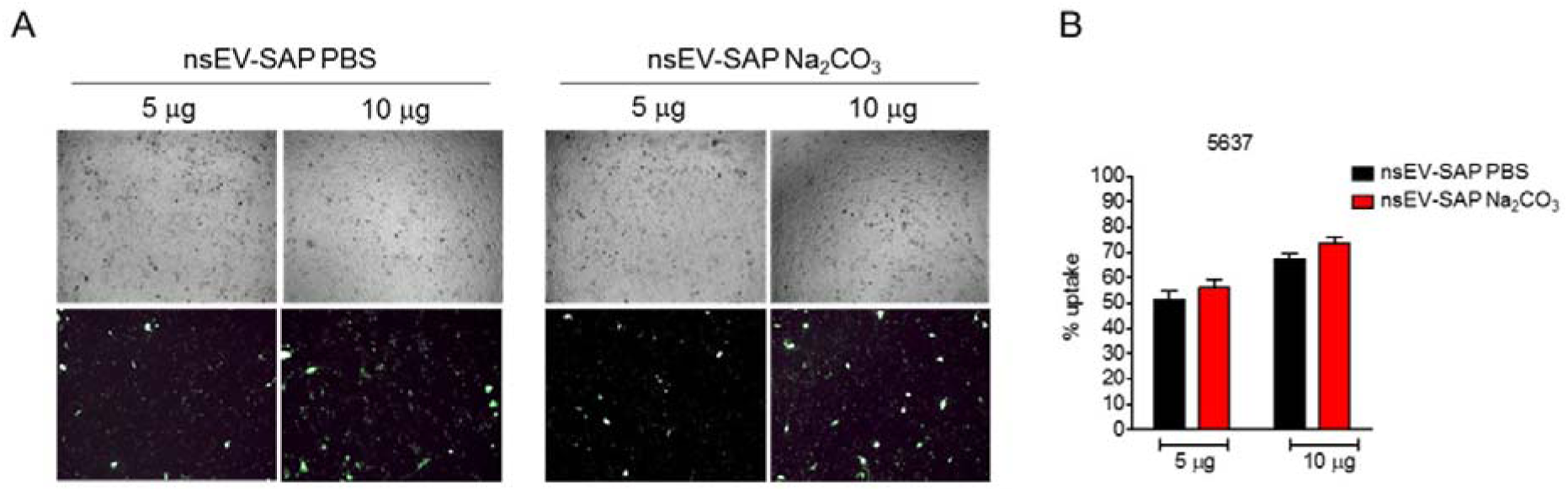
Evaluation of sodium carbonate-treated SAP-bearing nsEVs uptake. A. Bright field (upper row) and fluorescent (lower row) microscopy images of 5637 cells 24 hours after incubation with HEK293-derived nsEVs treated with phosphate buffer saline (PBS, left) or sodium carbonate (Na_2_CO_3_, right) in the presence of SAP and stained with Vybrant Dio lipophilic dye. Different concentrations of nsEVs (5 and 10 µg) were used for comparison. Representative images of three independent experiments. B. Flow cytometry analysis of 5637 cells 24 hours after treatment with 5 or 10 µg of PBS or Na_2_CO_3_-treated SAP-bearing nsEVs. Results are shown as mean fluorescence percentage + SD from 3 independent experiments.

5637 bladder cancer cells were then treated for 72 hours with increasing doses of SAP-bearing HEK nsEVs (corresponding to 5, 10, and 20 µg of protein) obtained by incubation with the toxin after prior incubation with sodium carbonate or PBS, as previously described. Sodium carbonate-treated SAP-bearing nsEVs (termed nsEV-SAP Na_2_CO_3_) strikingly affected cell viability in a dose-dependent manner and to a significantly improved extent compared to the nsEVs recovered after PBS treatment (nsEV-SAP PBS). Indeed, it can be appreciated that already the lowest dose of nsEV-SAP Na_2_CO_3_ used (corresponding to 10^9^ particles, with an estimated SAP content of 500–750 ng, based on observed loading capacity, Figure 3B and C) is higher than the effective IC_50_ dose (a dose that kills over 50% of recipient cells) (Figure 5). This observation is consistent with successful intravesicular protection and intracellular delivery of SAP.

**Figure 5.**
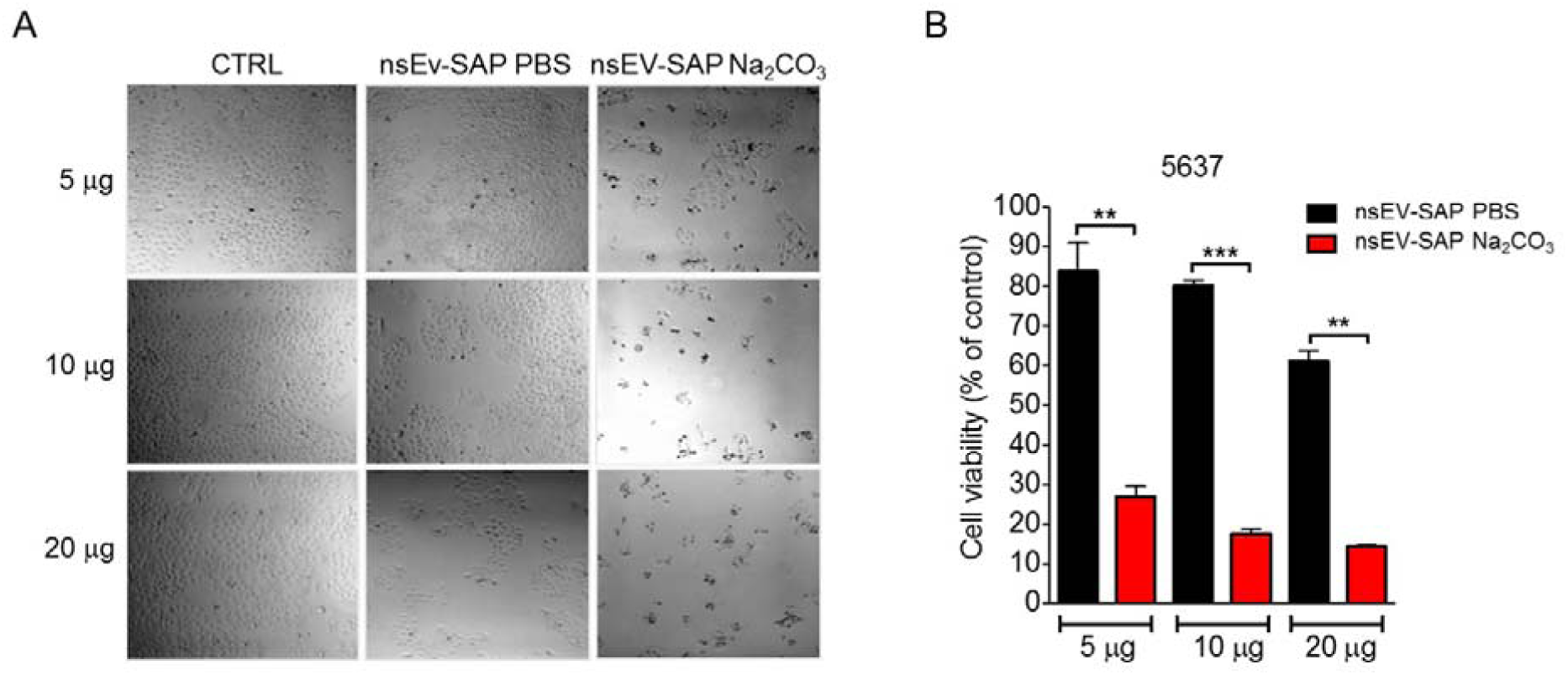
SAP-loaded nsEVs toxic activity evaluation on 5637 cancer cells. A. Bright field microscopy images of 5637 cells 72 hours after treatment with 5, 10 or 20 µg of SAP-bearing nsEVs (nsEVs-SAP) following sodium carbonate or PBS incubations. B MTT assay on 5637 cells after 72h incubation with 5, 10 or 20 μg of SAP-bearing nsEVs treated with sodium carbonate (red) or PBS (black). Results are shown as mean percentage residual viability to control + SD from three independent experiments. Test t ** *p* < 0.01, *** *p* < 0.001

We could also observe, although much smaller, a drop of cell viability upon treatment with a highest dose of nsEV-SAP PBS and hypothesized that this could be due to the saporin sticking to the nsEV surface. To test this hypothesis and reduce this surface association, accurately estimate the amount of encapsulated drug, and to guarantee limited potential by-stander toxic effects of the unincorporated fraction, we added sodium chloride washing steps following SAP encapsulation. As evidenced by western blot analysis (Figure 6A) of the processed nsEV samples, the SAP band was well detected in the carbonate-treated sample, whereas no SAP signal was retained in the PBS-treated control. By comparing the intensity of the nsEV-SAP band to that of serial dilutions of free protein, we estimated that 10 μg of loaded nsEVs contained approximately 1 μg SAP, which is in line with the loading efficiency estimated using fluorescently labelled protein in a prior experiment. At the same time, our hypothesis that some saporins bind to the nsEV surface was confirmed by the fact that the addition of sodium chloride washing resulted in lower cytotoxicity of nsEV-SAP Na_2_CO_3_ on tumor cells (Figure 5B) with an effective toxic dose, measured as IC_50,_ corresponding to more than 10 µg of nsEVs bearing SAP for washed samples, with respect to less than 5 µg of unwashed nsEVs used in a prior experiment (Figure 5A).

**Figure 6.**
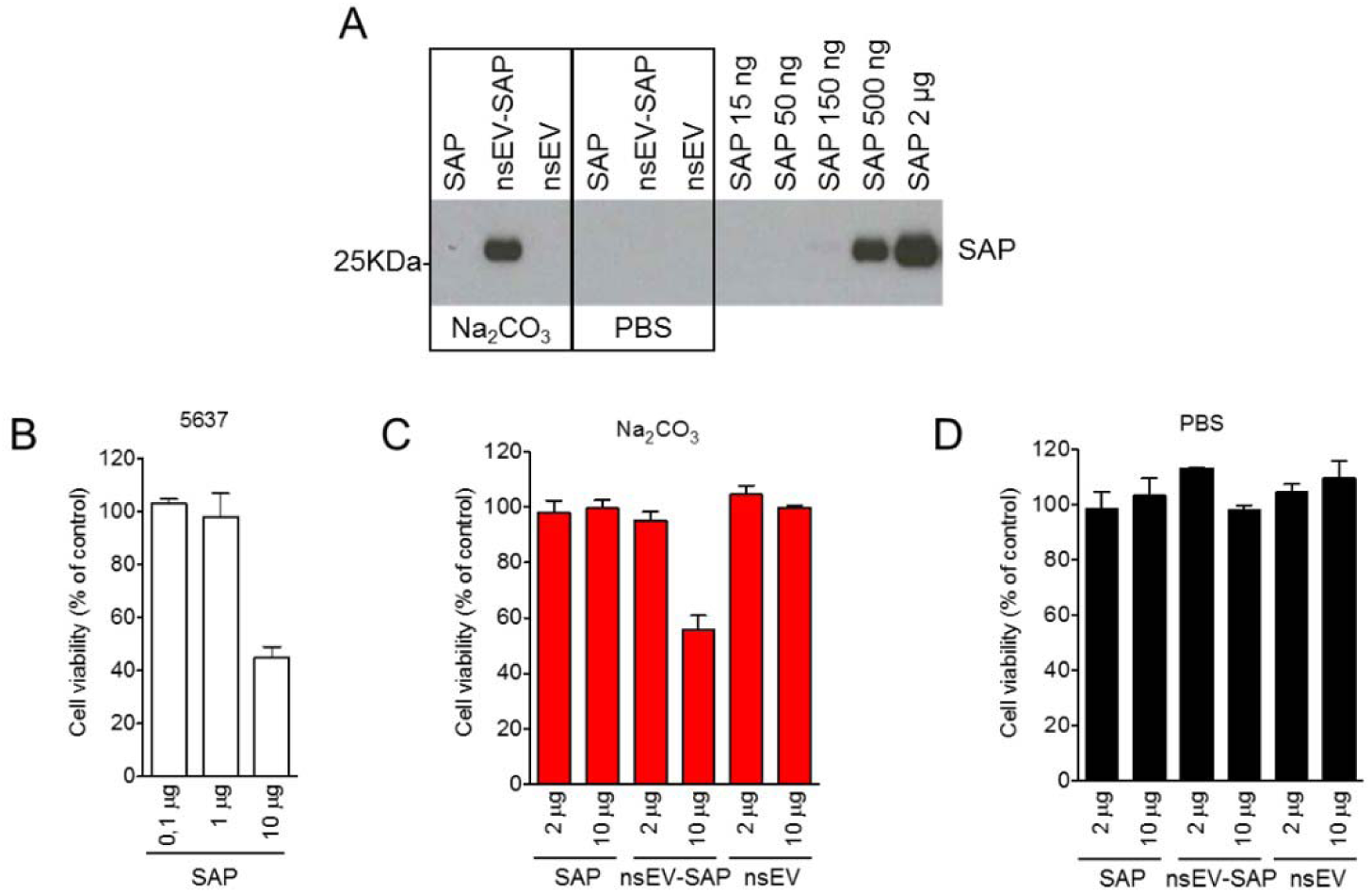
Elimination of SAP excess from nsEVs surface and evaluation of sodium carbonate encapsulation efficiency. A. Western blot analysis of SAP from sodium carbonate (Na_2_CO_3_) or phosphate buffer saline (PBS)-treated nsEVs. Excess of SAP was removed by washing steps with sodium chloride. 5 µg of nsEV proteins are loaded per lane. Non-loaded nsEVs and increasing concentrations of SAP alone were used as control. B. MTT assay on 5637 bladder cancer cells after 72 hours incubation with scalar concentrations of SAP. MTT assay on 5637 bladder cancer cells after 72 hours incubation with 2 and 10 µg of SAP, SAP-bearing nsEVs (nsEV-SAP) or nsEVs alone after Na_2_CO_3_ (C) or PBS (D) treatments. Samples were filtered with 100 kDa MWKO centrifugal devices to remove any unincorporated SAP. Results are shown as mean percentage residual viability to control + SD from three independent experiments.

Notably, we demonstrated that a conspicuously higher concentration of SAP toxin alone (approximately 8 μg of free toxin with respect to 1 μg of nsEV-encapsulated toxin) was needed to reach the IC_50_ under the same conditions (Figure 6B and C). This is likely due to the easier internalization of nsEV-delivered SAP than that of toxin alone. Furthermore, the toxic activity on cancer cells could not be influenced by the presence of the buffer, nor by the presence of the excess of non-encapsulated SAP, as both of them are completely eliminated during the washing procedure with centrifugal filter columns (Figure 6C and D).

## Discussion

In the last decades, the use of EVs as drug nanocarriers has revealed as a challenge for many research groups. Owing to their natural origin and morphological characteristics, the biomedical applications of these vesicles can overcome many of the drawbacks faced by artificial nanoparticles. Many strategies have been explored to either actively or passively load exogenous therapeutic cargoes into EVs, including genetic engineering of parental cells or direct application of mechano-physical stress to purified vesicles to perturb membrane integrity.^6^ One of the most effective and commonly employed approaches is electroporation, which has been widely used by many research groups to incorporate a wide range of molecules, including siRNAs, small molecules, and enzymes.^4,6,7,30^ However, when used to load exogenous molecules into EV lumen, it results in a very low loading efficiency (0.2% in the case of SAP), accompanied by a severe structural alteration of EV morphology.^6,15^ Overall, the exposure of vesicles to an electric field strongly affects their properties, causing aggregation and damage with a significant loss of the starting material.

In the present study, we took advantage of a highly concentrated alkaline buffer, sodium carbonate, to establish a method that, throughout pH alteration, can reversibly perturb nsEV membranes and favor the encapsulation of exogenous molecules. Sodium carbonate was first used in 1982 by Fujiki *et al*. to isolate intracellular/organellar membranes.^31^ They demonstrated that incubation of purified fractions of rat liver endoplasmic reticulum, peroxisomes, or mitochondria in sodium carbonate induced the conversion of membranes into membrane sheets, which retained only integral membrane proteins, but not lipid-anchored proteins. In fact, alkaline pH decreases noncovalent protein–protein interactions, releasing weakly attached peripheral membrane proteins without affecting membrane integrity. Here, we first proved that the use of a high concentration of sodium carbonate (1M) can preserve the morphological and biochemical properties of vesicles. For this purpose, we combined several assays to assess the independent vesicle parameters that are indicative of their amount and integrity. The decrease in particle count and protein content to 85–65% respectively, with respect to PBS treated control indicated a moderate loss of nsEVs due to the loading protocol. However, it is possible that this loss is overestimated, as alkaline washing is likely to clear up the EV sample of associated proteins and decrease the amount of protein aggregates that may appear as particles. If we use the lipid content as an indicator of the amount of EVs in our preparation, the loss of nsEVs is negligible. Therefore, it is likely that this alkaline wash adds value to the purity of EV preparations. Moreover, the preserved integrity of nsEVs recovered after carbonate treatment is supported not only by the well-maintained protein or lipid content per particle but also by the preserved recognition of surface proteins and glycans in ELISA and lectin binding assays. Therefore, rather than interfering with the vesicle corona, this treatment ensured the fine display and functionality of the EV membrane domains, potentially mediating recognition and uptake. Finally, the fully preserved functionality of the recombinant protein (CD9-RFP fusion) in engineered HEK nsEVs upon carbonate buffer incubation is a fine proof that this loading strategy might be applied to EVs with modified membrane domains, and thus, is compliant and complementary with other strategies for tailored nsEV fabrication.

We envisaged that carbonate treatment would prompt exogenous molecule encapsulation by co-incubation depending on the type and size of the molecule used. This hypothesis was confirmed by the observation that molecules with different structural properties (dextran and SAP) are incorporated in a dose-dependent manner. However, the distinctive steric hindrance of the two molecules was reflected in the encapsulation efficiency, which was higher when globular enzyme SAP was used. Nonetheless, we showed that the sodium carbonate-induced loading capacity appears to be significantly improved compared to electroporation under our experimental conditions. Importantly, we excluded SAP aggregation on the surface by introduction of washing with saline solution. The retention of the SAP signal in carbonate buffer-treated nsEVs, but not in PBS-incubated nsEVs, clearly indicated that a significant fraction of endogenous protein was indeed encapsulated and protected by the vesicular membrane. The difference between sodium carbonate-treated SAP-bearing nsEVs and PBS control treatment was unambiguously evident in the *in vitro* cytotoxic assay, where the former was able to strongly affect cell viability at a very low dose. Conversely, PBS-treated SAP-bearing nsEVs did not produce any marked toxic effects even at the highest concentration used. It is worth mentioning that in our study, an effective *in vitro* dose of SAP-loaded nsEVs corresponded to 10^9^–10^10^ vesicles, being thus in a reported dose range for EV based functional assays^32^ and bearing an effective concentration of SAP to cause striking, tumor cell-killing, activity. Importantly, the effective SAP IC_50_ was in the nanomolar range, a dose significantly lower than the necessary IC_50_ of a free toxin and comparable to the efficacy of targeted SAP variants reported in the literature.^28,29^ Finally, we demonstrated that the use of a physiological concentration of sodium chloride in the washing step allows the surface-attached portion of SAP to be released, lowering the weak protein interactions with the vesicle membrane. Thus, the biological effect is confined to the encapsulated toxin, rendering its delivery to recipient cells safer and more controlled. We could observe that, while for 75 kDa dextran the efficiency of carbonate buffer treatment resembles that of electroporation (around 0.2–0.3%), its efficiency in incorporation of 25 kDa globular protein reaches 10–15%, which is about 10 times higher that of the commonly used electroporation protocol (the efficiency of loading by electroporation in our hands was 1–1.5 %, higher than that typically reported in other studies).^15^ Our results also showed that pre-treatment of EVs with carbonate buffer increased the loading efficiency of the protein payload compared to that reported for passive incubation in other studies. For instance, the efficiency of loading of the antioxidant enzyme catalase into macrophage EVs via passive incubation was reported to be <5%.^6^ The addition of membrane permeabilization agents, such as detergents (i.e., saponin), has been proposed to increase the efficacy of passive incubation loading strategies, but they pose additional downstream steps and problems.^33^ They can hamper the downstream analytics or reactivity of EV surface proteins and necessitate additional time-consuming steps that augment material loss and decrease reproducibility. Our method is fully reversible, does not leave any residual additives or reactives, and fully preserves EV structure and function.

In conclusion, we demonstrated that pH alteration induced by high concentrations of sodium carbonate is a safe and effective method for arranging nsEVs as drug nanocarriers, preserving their size and structure, allowing conspicuous loading of a therapeutic protein, and mediating the desired biological outcome in recipient cells. Further investigations will confirm the transferability of our method to a broad spectrum of EV sources. Nevertheless, to achieve the effective intracellular delivery of EV contents, future experiments will be performed in order to assess the compatibility of this loading protocol with preserved functionality of different types of exogeneous cargo of interest and, on the other side, to stably customize EV membrane with chemically conjugated targeting moieties, which must enhance a selective and preferential cellular uptake by target cells exploiting a receptor-mediated endocytosis uptake rather than micropinocytosis.

## Disclosure

The authors report no conflicts of interest in this work

## Acknowledgments

Part of this work was conducted at the Advanced Light and Electron Microscopy BioImaging Center (ALEMBIC) of the San Raffaele Scientific Institute and Vita-Salute University.

