## Supplementary Figures for "Novel loading protocol combines highly efficient encapsulation of exogenous therapeutic toxin with preservation of extracellular vesicles properties, uptake and cargo activity"

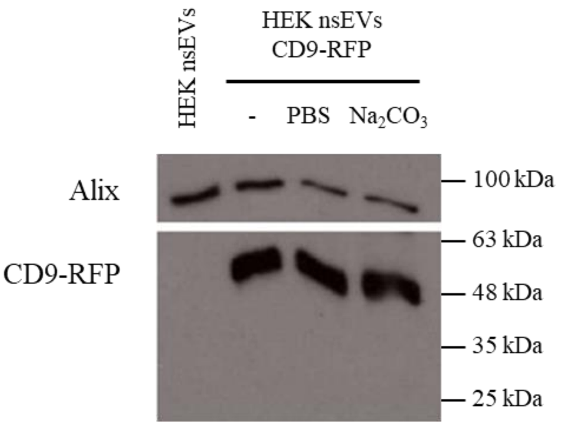


**Fig. S1. HEK293 derived nsEVs structural stability upon sodium carbonate treatment.** Western blot analysis Alix and RFP from untreated (-), PBS treated and sodium carbonate ($Na_2CO_3$) treated nsEVs derived from CD9-RFP expressing HEK293 cells. HEK293 WT derived nsEVs were used as negative control for CD9-RFP expression. CD9-RFP has been detected using anti-RFP antibody.

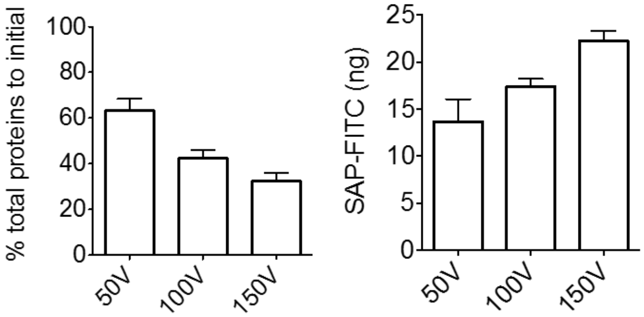


**Fig. S2. Protein recovery and encapsulation efficiency evaluation after increasing voltages exposition and higher nsEVs:SAP-FITC ratios.** A. BCA assay for quantification of total protein recovery after co-incubation of HEK293-derived nsEVs (25 ug/ml) with SAP-FITC (1:2 molecular ratio) and exposition to 50, 100, 150 V electroporation (see material and methods). Results are shown as percentage to initial. B. Quantification of SAP-FITC incorporation after incubation with HEK293-derived nsEVs (25 ug/ml) at a 1:2 molecular ratio and 50, 100, 150 V electroporation (see Material and Methods) by spectrofluorometer analysis.
